# Nucleotide GPT: Sequence-Based Deep Learning Prediction of Nuclear Subcompartment-Associated Genome Architecture

**DOI:** 10.1101/2024.11.27.625761

**Authors:** Shae M. Mclaughlin, Sajad H. Ahanger, Daniel A. Lim

## Abstract

The spatial organization of the genome within the nucleus is partially determined by its interactions with distinct nuclear subcompartments, such as the nuclear lamina and nuclear speckles, which play key roles in gene regulation during development. However, whether these genome-nuclear subcompartment interactions are encoded in the underlying DNA sequence remains poorly understood. The mechanisms for gene regulation are primarily encoded in noncoding DNA sequences, but deciphering how these sequence features control gene expression remains a significant challenge in genomics. Here, we present Nucleotide GPT, a transformer-based model that predicts genomic associations with spatially distinct, physical nuclear subcompartments from DNA sequence alone. Pre-trained on a diverse set of multi-species genomes, we demonstrate Nucleotide GPT’s genomic understanding through evaluation on diverse prediction tasks, including histone modifications, promoter detection, and transcription factor binding sites. When finetuned to predict genome interactions with two separate nuclear subcompartments – the lamina of the inner nuclear membrane and nuclear speckles that lie more interior – Nucleotide GPT achieves an average accuracy of 73.6% for lamina-associated domains (LADs) and 79.4% accuracy for speckle-associated domains (SPADs), averaged across three cortical development cell types. Analysis of the model’s learned representations through Uniform Manifold Approximation and Projection (UMAP) reveals that Nucleotide GPT develops internal embeddings that effectively distinguish LADs from inter-LADs, with predicted probabilities closely corresponding to experimentally determined LAD classifications. When examining these representations in the context of cell type-invariant constitutive LADs (cLADs) compared to cell type-specific LADs, the model assigns lower confidence scores to cell type-specific LADs compared to cLADs that are conserved across neuronal differentiation, suggesting sequence features may play a stronger role in maintaining cLAD associations. Examination of the model’s attention patterns at correctly classified regions suggests that specific sequence elements govern model decision making about nuclear subcompartment associations. Our results demonstrate the utility of transformer architectures for studying three-dimensional (3D) genome organization and substantiate a role for DNA sequence in determining nuclear subcompartment associations.

## 1 Introduction

The three-dimensional (3D) organization of chromatin within the cell nucleus is known to play a critical role in the regulation of gene expression (1). The cell nucleus contains distinct subnuclear structures that play key roles in genome organization and gene regulation. The nuclear lamina, a fibrous protein network lining the inner nuclear membrane, tethers chromatin to the nuclear periphery through interactions with large chromatin domains called lamina-associated domains (LADs). These LADs are generally characterized by low gene density and transcriptional repression (2). Another prominent nuclear structure is the nuclear speckle, a membraneless organelle enriched in pre-mRNA splicing factors and various proteins involved in gene expression. Speckle-associated domains (SPADs) are less well-known, but genes within this subcompartment are highly expressed and exhibit greater splice isoform diversity (3). Although the importance of these nuclear subcompartments in genome organization is well established, whether there are specific DNA sequences that underlie LAD and SPAD formation remain poorly understood.

Large language models, also known as foundation models, present an opportunity to leverage the transformer architecture’s capacity for language modeling to unearth novel insights about the complex relationships and dependencies in DNA sequences, transcriptional regulation, and the functional effects of regulatory elements (4). This approach is particularly valuable for studying gene regulation, where there are clear parallels between natural language and regulatory DNA. Individual cis-regulatory elements (CREs) can exhibit different functions depending on their cellular context, and multiple CREs, even when separated by large genomic distances, can work together to regulate gene expression through alternative promoter usage. This behavior suggests the existence of polysemy and distant semantic relationships within genomic sequence, key properties of natural language (5).

The development of genomic foundation models has evolved rapidly in recent years. DNABERT was the first to show that transformer architectures, originally developed for processing human language, could be adapted to understand DNA sequences and their regulatory patterns (5). Foundation models for genomic sequences have proven effective across diverse biological prediction tasks, including promoter prediction (6) (7), gene expression prediction (8), DNA methylation prediction (9), chromatin state analysis (10), promoter-enhancer interaction prediction (11) (12), transcription factor (TF) binding prediction (13), variant effect prediction (14), and gene network prediction (15). The broad success of these models in capturing various aspects of genome function suggests they can extract biologically meaningful patterns from DNA sequence alone. While these foundation models have proven effective for many genomic prediction tasks, their application to epigenomic tasks, such as 3D genome organization, has been limited. Yang & Ma (16) demonstrated that transformer architectures could predict chromosome spatial positioning from sequence, using tyramide signal amplification sequencing (TSA-seq) data in cultured cells, but deep learning prediction of in vivo nuclear subcompartment associations remains unexplored.

Whether DNA sequence plays a significant role in determining nuclear subcompartment associations remains an open question in genome organization. Previous work from our lab using Genome Organization with CUT&RUN technology (GO-CaRT) enabled the mapping of LADs and SPADs with relatively small numbers of cells isolated from in vivo tissues (17). These studies demonstrated that LADs are characterized by distinct epigenetic signatures, including enrichment of histone lysine-9 dimethylation (H3K9me2) and moderate levels of histone H3 lysine-9 trimethylation (H3K9me3), while histone H3 lysine-27 trimethylation (H3K27me3) is predominantly localized to LAD borders and depleted within LADs. These findings highlight the importance of histone modifications in LAD organization but leave open the question of whether underlying DNA sequence contributes to these associations. The emergence of foundation models in genomics, with their ability to learn complex sequence patterns and be fine-tuned for specific tasks, presents an opportunity to test whether DNA sequence alone contains sufficient information to predict nuclear subcompartment associations.

Here, we introduce a new pre-trained decoder-only genome transformer called Nucleotide GPT, which is designed to predict genome association with nuclear subcompartments from DNA sequence alone. Pre-trained on reference genomes from multiple species including human, mouse, and macaque, the model develops broad genomic understanding before being fine-tuned to predict LADs and SPADs. Given that cell culture can impact genome architecture (17), we focused on three specific cell types acutely isolated from the developing human brain: radial glia (RG) – the primary neural precursor cells – which give rise to intermediate progenitor cells (IPC) that then produce postmitotic excitatory neurons (eN). Nucleotide GPT demonstrates how transformer models’ transfer learning capabilities can be extended to epigenomic prediction tasks, providing a computational framework for exploring sequence determinants of nuclear genome architecture.

## 2 Methods

### Model Architecture

Nucleotide GPT is a decoder-only model which employs a transfer learning approach through a two-stage paradigm of pre-training and fine-tuning. During pre-training, it develops broad genomic sequence understanding by learning from reference genomes across multiple species through self-supervised causal language modeling. In fine-tuning, these foundational capabilities are adapted to classify nuclear-associated genome organization states by retraining the model’s parameters on sequences experimentally associated with nuclear lamina or nuclear speckles. This fine-tuning process enables efficient transfer of cross-species genomic knowledge to the specific task of predicting aspects of 3D genome organization, requiring minimal architectural modifications to the base model.

The model’s architecture is adapted from Vaswani et al. (18). At the input layer, DNA sequences are tokenized and embedded. To encode positional information, we use rotary positional embeddings (RoPE) (19). RoPE is applied to the input embeddings as follows:

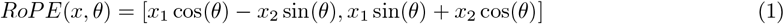

where *x*_1_ and *x*_2_ are the first and second half of the embedding vector, and *θ* is a function of the position and dimension.

The core of our model consists of multiple transformer layers. Each layer implements a multi-head attention mechanism, followed by a position-wise feed-forward network. The multi-head attention operation is defined as:

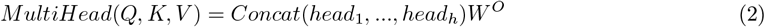

where:

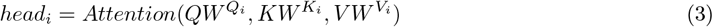

The attention function uses scaled dot-product attention:

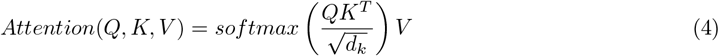

We implement this attention computation using Flash Attention (20), an IO-aware algorithm that enables efficient training while maintaining accurate attention calculations. Following the attention mechanism, each transformer layer applies a position-wise feed-forward network:

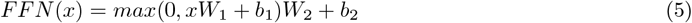

Instead of the traditional layer normalization, we employ root mean square (RMS) layer normalization for improved stability (21):

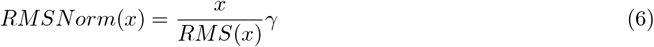

where:

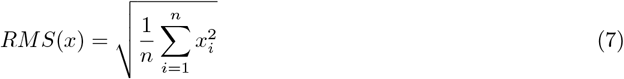

The output of each transformer layer can be summarized as:

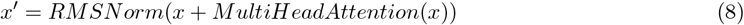

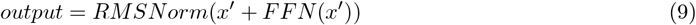

The final layer of the model projects the transformer output to predict the association probabilities for nuclear lamina and nuclear speckles.

We use a cross-entropy loss function for training:

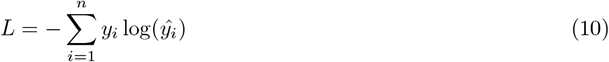

where *y*_*i*_ are the true labels and *ŷ*_*i*_ are the predicted probabilities.

The model is implemented using JAX libraries for efficient computation and easy deployment on TPU hardware. We employ a custom training loop that includes learning rate scheduling, gradient clipping, and AdamW optimization. Pre-training was conducted on four TPUs for approximately one day.

### Model Training

Nucleotide GPT is pre-trained on a cross-species dataset that includes the human reference genome, the Mus musculus genome, the Macaca mulatta genome, the Danio rerio genome, and the Drosophila melanogaster genome. Unlike previous genome language models such as DNABERT which use k-mer tokenization (where each token represents a sequence of k nucleotides), our model employs a simple nucleotide-level tokenization scheme. Each nucleotide (A, T, G, C) is represented as an individual token in the vocabulary, along with special tokens for padding, and unknown bases. This approach preserves the biological integrity of the 8kb non-overlapping sequences the model is trained on, avoiding arbitrary fragmentation that could disrupt significant motifs and regulatory elements.

### Evaluation

Nucleotide GPT is fine-tuned and evaluated on the Genome Understanding Evaluation (GUE) benchmark (4) and compared against leading genomic foundation models including HyenaDNA (1K version) (22), Nucleotide Transformer variants (NT-500M-1000g, NT-2500-multi, NT-50M-multi-V2) (23), DNABERT-2 (4), and dnaGrinder (24). The model’s performance was assessed using published results from the dnaGrinder benchmark evaluation across multiple genomic prediction tasks including promoter detection, splice site prediction, transcription factor binding site (TFBS) prediction, and epigenetic marks prediction. The COVID variant prediction task was excluded as the model was not trained on viral DNA sequences. After single-epoch fine-tuning on the GUE training dataset, performance was assessed on a held-out test set, using data obtained from the DNABERT-2 GitHub repository.

### Model Interpretability

To analyze how the model represents genomic sequences, we employed two complementary approaches: UMAP visualization of sequence embeddings and analysis of attention scores. For UMAP visualization, we extracted embeddings from the model’s final layer after fine-tuning and performed dimensionality reduction using the UMAP algorithm (25), with default parameters. This allowed us to visualize the organization of sequence representations in two dimensions and examine how they correspond to biological annotations including nuclear lamina and nuclear speckle association.

To interpret which genomic regions were most influential for the model’s predictions, we developed a method to analyze the transformer’s attention patterns. For unseen examples held out during fine-tuning for use in later validation, we extracted attention scores from all layers and attention heads during the model’s forward pass. These attention matrices (12 layers × 8 heads × 8192 × 8192) were summed across layers and heads, then summed again along one dimension to obtain a single attention score per nucleotide position. This aggregation strategy captures the cumulative importance of each position across all model components. To identify significant attention peaks, we implemented a sliding window approach that compares local attention intensities to background levels. We calculated z-scores using a rolling mean and standard deviation (window size = 250 bp), with regions exceeding a threshold of z > 3 identified as attention spikes. To avoid redundant peak calling, peaks within 200 bp of each other were merged, with the highest scoring peak retained. Edge effects were controlled for by trimming 500 bp from the start and 200 bp from the end of each sequence. The resulting attention profiles were converted to bedGraph format for visualization in the UCSC Genome Browser, enabling direct comparison between model attention patterns and known genomic features including GENCODE V47 gene models (26) showing transcript structure including exons, introns and transcription start sites, regulatory elements from ENCODE (27) and ORegAnno (28) databases, ReMap ChIP-seq signals (29), Hi-C chromatin interaction data (30), conservation scores (31), predicted CpG islands (32), and predicted transcription factor binding sites from JASPAR 2024 (33). This approach provides a method for interpreting transformer model behavior in the context of genomic sequence analysis, potentially revealing previously unrecognized sequence elements important for nuclear organization.

### Data Collection and Preprocessing

Human brain tissue samples (GW17) were collected from de-identified donors with previous patient consent in strict observance of the legal and institutional ethical regulations. All protocols were approved by the Human Gamete, Embryo and Stem Cell Research Committee (GESCRC) and the Institutional Review Board (IRB) at the University of California, San Francisco (UCSF). Genome Organization with CUT&RUN technology (GO-CaRT), and fluorescence-activated nuclei sorting (FANS) was performed as previously described (34).

Sequencing reads were first trimmed to remove adapters and low-quality sequences using Trim Galore with default parameters. The trimmed reads were then mapped to the human genome (hg38) using Bowtie2. PCR duplicates were removed using Picard tools. Properly paired reads with high mapping quality (MAPQ score > 30) were kept for further analysis. For domain calling, the genome was binarized into 10kb bins and LADs and SPADs were called using SEACR (v1.3) that is specifically designed to call broad enriched regions as per instruction. Each target data bedgraph was normalized by the corresponding IgG control, where the peak of the curve was set as the threshold during the domain calling.

To preprocess the LADs and SPADs for fine-tuning, the human genome was divided into 8kb bins. Bins with significant or complete overlap with LADs and SPADs were classed as ‘LAD’ or ‘SPAD’. Bins with no overlap were classed as ‘inter-LAD’ or ‘inter-SPAD’. Bins that contained a LAD or SPAD border were classed as ‘LAD boundary’ or ‘SPAD boundary’.

## 3 Results

To investigate the sequence determinants of nuclear organization, we developed Nucleotide GPT, a transformer-based decoder-only model that processes DNA sequences at single-nucleotide resolution. The model was pretrained on reference genomes from multiple species including human, mouse, macaque, zebrafish, and fruit fly to develop broad genomic understanding. Unlike previous approaches that use k-mer based tokenization, we implemented single-nucleotide tokenization to preserve the complete linear organization of DNA sequences. The pre-trained model was then fine-tuned to predict nuclear subcompartment associations using experimentally determined LADs and SPADs from three cell types of the developing human cortex: RG, IPC, and eN. We first evaluated the model’s general genomic understanding through a suite of prediction tasks before assessing its ability to capture nuclear organization patterns.

### Nucleotide GPT Achieves Comparable Performance to State-of-the-Art Models

To evaluate the model on promoter detection, two tasks were used: core promoter detection using 70 bp sequences (−34 to +35 bp around transcription start site (TSS)) (**Fig. 1a**) and standard promoter detection using 300 bp sequences (−249 to +50 bp around the TSS) (**Fig. 1b**). On standard promoter detection, Nucleotide GPT achieved strong performance on non-TATA promoters (95.57%), approaching state-of-the-art accuracy (NT-2500-multi: 97.11%), while showing lower performance on TATA-box containing promoters (78.30% vs NT-50M-multi-V2’s 93.63%). This pattern persisted in core promoter detection, with Nucleotide GPT achieving state-of-the-art performance on non-TATA promoters (83.78%) and reduced accuracy on TATA-containing sequences (73.25% vs NT-50M-multi-V2’s 89.06%). These results suggest that while Nucleotide GPT effectively captures the broader regulatory context of promoter regions, it may be less efficient at identifying specific core promoter sequence motifs, particularly the TATA-box element.

**Figure 1:**
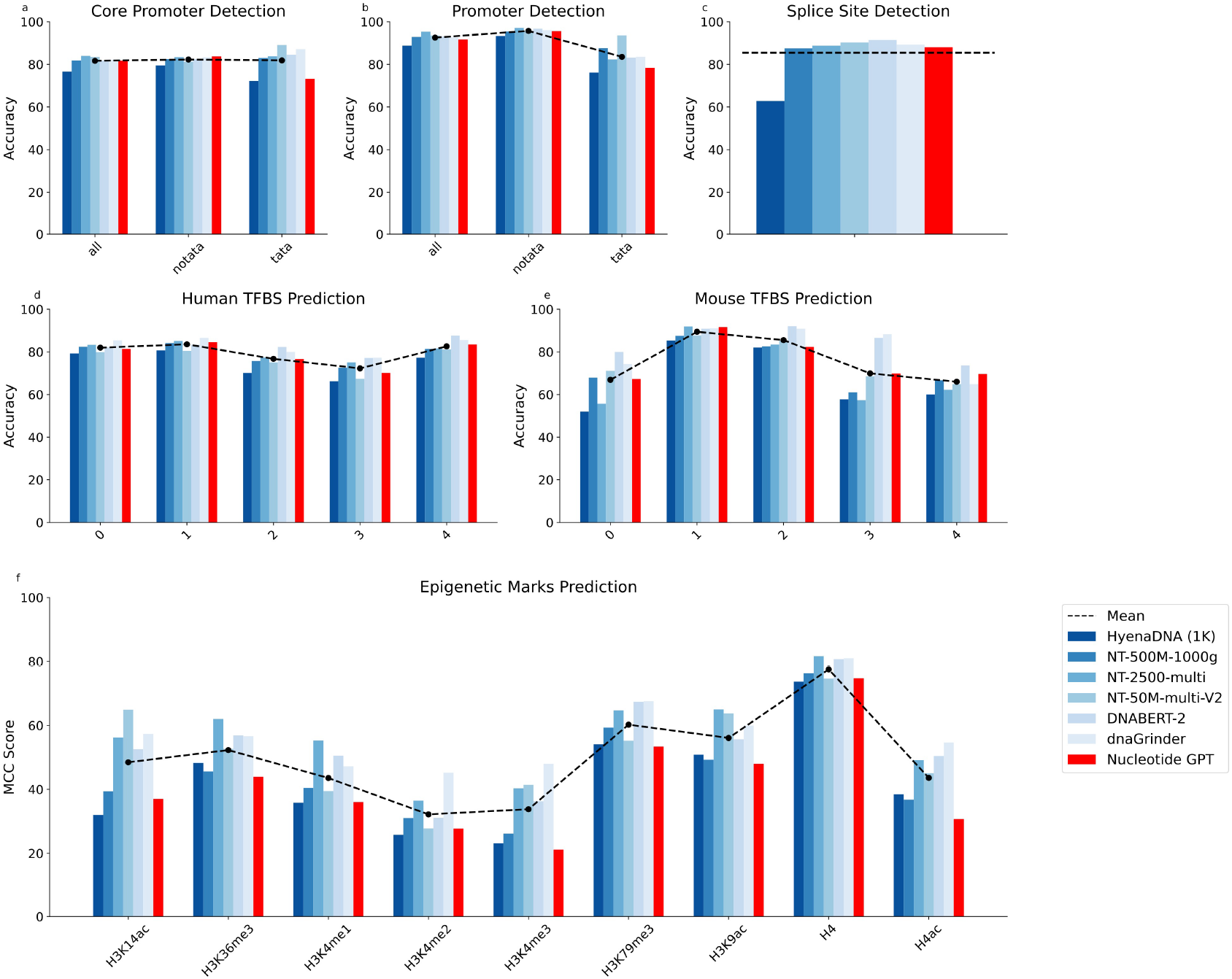
Genome Understanding Evaluation (GUE) performance comparison of Nucleotide GPT against existing genome foundation models. Comparison of Nucleotide GPT (red) with existing models (blue shades) across existing genome foundation models. (a) Core promoter detection (70 bp) accuracy across all promoters, TATA-containing promoters, and TATA-less promoters. (b) Promoter detection (300 bp) accuracy across all promoters, TATA-containing promoters, and TATA-less promoters. (d) Human TFBS prediction accuracy for five different types of transcription factors (0-4). (e) Mouse TFBS prediction accuracy for the same five transcription factor types. (f) Performance on epigenetic mark prediction tasks across nine different histone modifications, measured by Matthews Correlation Coefficient (MCC).

We evaluated Nucleotide GPT’s performance on splice site prediction using 400 bp sequences from the human genome (**Fig. 1c**). Nucleotide GPT achieved 88.01% accuracy, comparable to DNABERT-2 (90.31%) and NT-50M-multi-V2 (87.52%), while substantially outperforming NT-500M-1000g (62.84%). This performance demonstrates Nucleotide GPT’s ability to identify complex splice site signals that are crucial for understanding alternative splicing mechanisms.

We evaluated Nucleotide GPT’s performance on TFBS prediction using 101 bp sequences in both human and mouse genomes. On human TFBS prediction, Nucleotide GPT maintained competitive performance across five different transcription factors, achieving 84.60% accuracy on TF-1 prediction, comparable to the state-of-the-art performance of NT-2500-multi (85.10%) (**Fig. 1d**). Performance on mouse TFBS prediction was particularly strong for certain factors, with Nucleotide GPT achieving 91.59% accuracy on TF-1, approaching NT-2500-multi’s leading performance (91.91%) and surpassing DNABERT-2 (85.86%) (**Fig. 1e**). While Nucleotide GPT showed lower performance on some factors (e.g., 70.10% vs DNABERT-2’s 77.20% on human TF-3), the differences were generally modest, demonstrating that Nucleotide GPT can effectively identify regulatory binding motifs.

On histone mark prediction tasks, while Nucleotide GPT generally performed below other models, the performance differences were modest (**Fig. 1f**). Across all ten histone mark prediction tasks, Nucleotide GPT performed, on average, 3.2 percentage points lower than NT-500M-1000g (95% CI: 1.8, 4.6), 14.5 percentage points lower than NT-2500-multi (95% CI: 10.8, 18.2), 8.9 percentage points lower than NT-50M-multi-V2 (95% CI: 1.7, 16.2), 11.5 percentage points lower than DNABERT-2 (95% CI: 8.0, 15.1), and 15.3 percentage points lower than dnaGrinder (95% CI: 11.2, 19.5). Collectively, these evaluations demonstrate that Nucleotide GPT achieves performance comparable to state-of-the-art genomic language models across diverse prediction tasks, suggesting the model develops meaningful internal representations of genomic sequence patterns.

### Nucleotide GPT Captures Nuclear Lamina Association Patterns

We fine-tuned Nucleotide GPT to classify sequences associated with LADs, inter-LADs, and LAD boundaries in three key neural cell types involved in cortical development: RG, IPC, and eN. The goal was to assess the pre-trained model’s ability to learn LAD associations based solely on DNA sequences in different stages of neuronal differentiation. For the RG sample, the model achieved 77.5% accuracy after 9,000 training steps. For the IPC sample, the model reached an accuracy of 74.5%. For the eN sample, the model achieved 68.9% accuracy. **Fig. 2** shows the training accuracy over the 9,000 steps for each cell type. The model’s performance varies across cell types, with the highest accuracy in RG and progressively lower accuracies in IPC and eN. This trend suggests that the sequence determinants of LAD associations may differ among these cell types or that the complexity of LAD organization increases during differentiation. We also fine-tuned Nucleotide GPT on a combined dataset encompassing all three cell types to create a cross-cell type model. This model achieved an overall accuracy of 71.3% on LAD classification.

**Figure 2:**
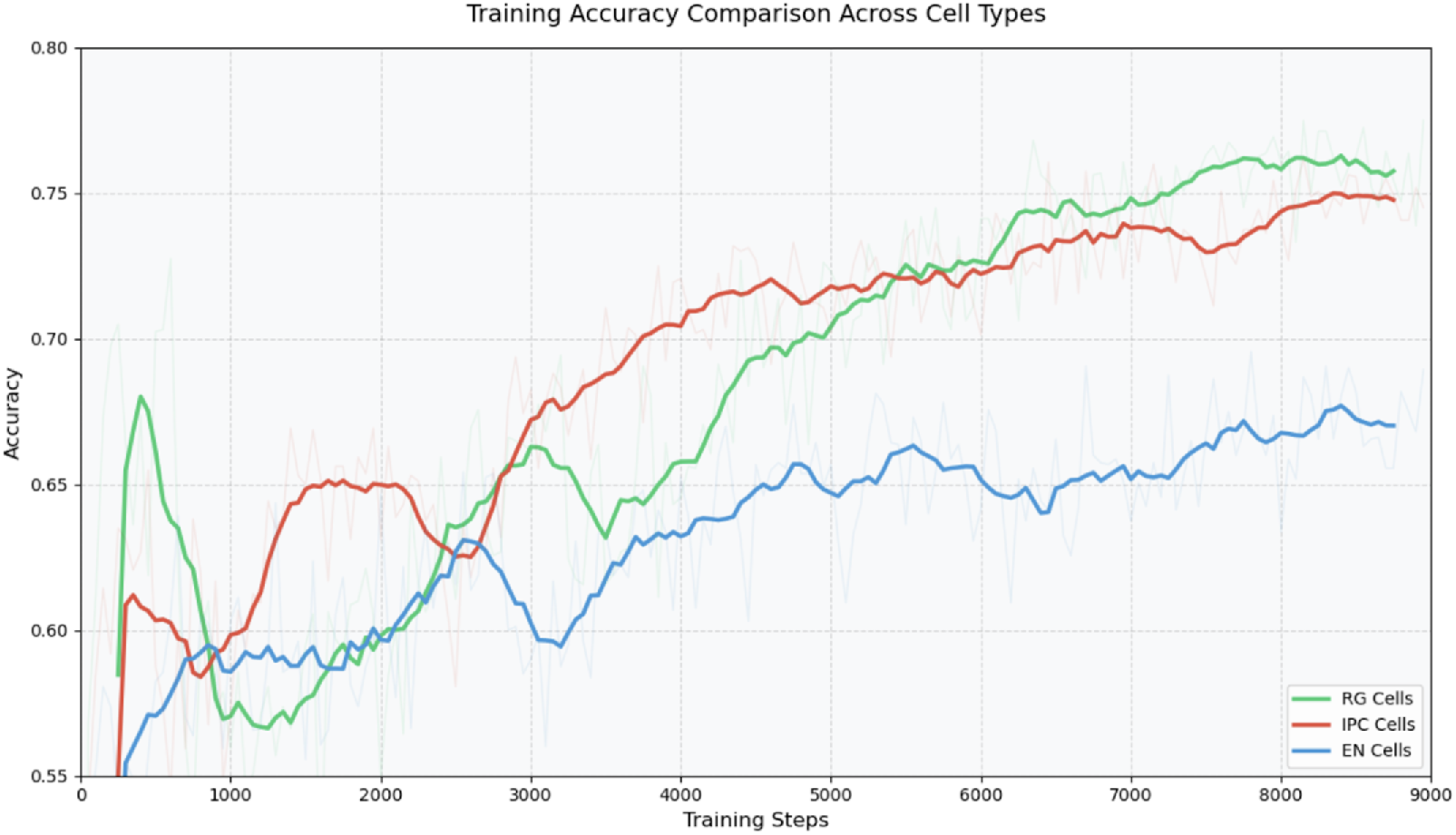
LAD classification fine-tuning training accuracy across three cortical development cell types. Training accuracy trajectories for lamina-associated domain (LAD) classification in three neural cell types: RG, IPC and EN. Each line represents the classification accuracy over training steps for a cell type-specific fine-tuning model, showing how well each model learns to identify LAD regions in the respective cell types.

To further understand the model’s predictive behavior, we analyzed its classification confidence across different categories of genomic regions (**Fig. 3**). The model showed significantly higher confidence in classifying conserved LADs (mean confidence = 0.540 ± 0.222) compared to RG-specific LADs (mean confidence = 0.476 ± 0.227). The model also demonstrated very high confidence in identifying inter-LADs, suggesting that these regions contain sequence features learned by the model that distinguish them from LAD regions. This analysis reveals that while the model achieves moderate accuracy in overall LAD classification, its confidence varies systematically with LAD conservation status, with the highest confidence for genomic regions that maintain consistent nuclear compartment associations across cell types.

**Figure 3:**
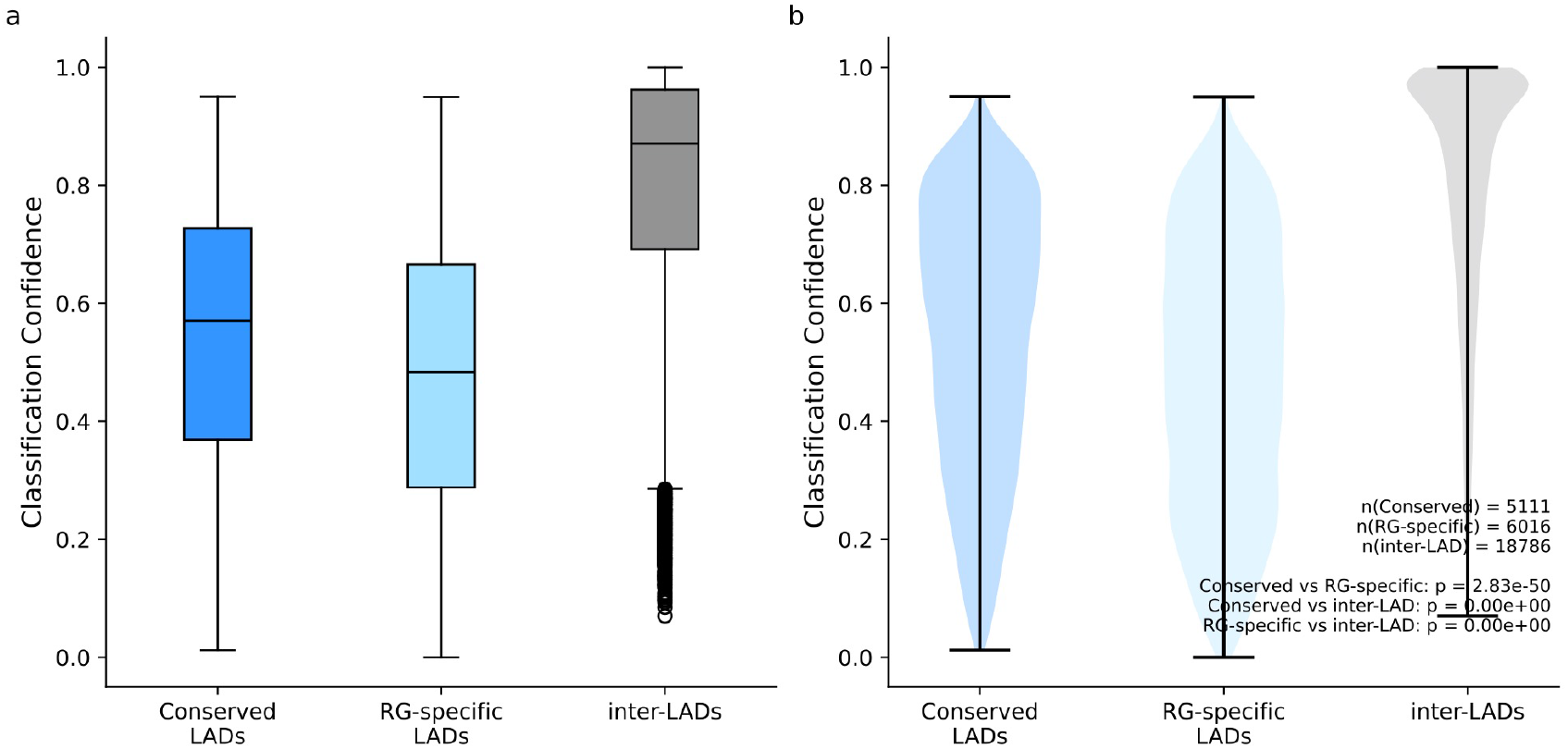
LAD classification confidence by conservation status. The model demonstrates varying levels of classification confidence across different categories of genomic regions. Box plots (a) and violin plots (b) show the distribution of classification confidence scores for conserved LADs (present in both RG and eN), RG-specific LAD (present only in RG), and inter-LADs (regions between LADs). Conserved LADs show significantly higher classification confidence (mean = 0.540 ± 0.222) compared to RG-specific LADs (mean = 0.476 ± 0.227), suggesting that sequence features may play a stronger role in determining constitutive LAD associations. Inter-LADs also show high classification confidence, indicating distinct sequence characteristics that distinguish them from LAD regions. Box plots show median (black line), interquartile range (box), and full range excluding outliers (whiskers). Violin plots illustrate the full distribution of confidence scores. n(Conserved) = 5,111; n(RG-specific) = 6,016; n(inter-LAD) = 18,786; all pairwise comparisons p < 2.83e-50.

To understand how the model represents these genomic sequences after fine-tuning, we analyzed the organization of the sequence embeddings using Uniform Manifold Approximation and Projection (UMAP) dimensionality reduction. The UMAP visualizations demonstrate a notable correspondence between the model’s predicted LAD probabilities and the actual LAD annotations (**Fig. 4a, b**). While there are no sharp boundaries or distinct clusters, regions annotated as LADs (**Fig. 4a**) correspond well to regions where the model predicts high LAD probabilities (**Fig. 4b**), and similarly, annotated inter-LADs (**Fig. 4a**) align with regions of low predicted LAD probability (**Fig. 4b**). This continuous gradient of probabilities in the embedding space suggests the model has learned to capture a spectrum of nuclear lamina association strengths rather than making binary classifications.

**Figure 4:**
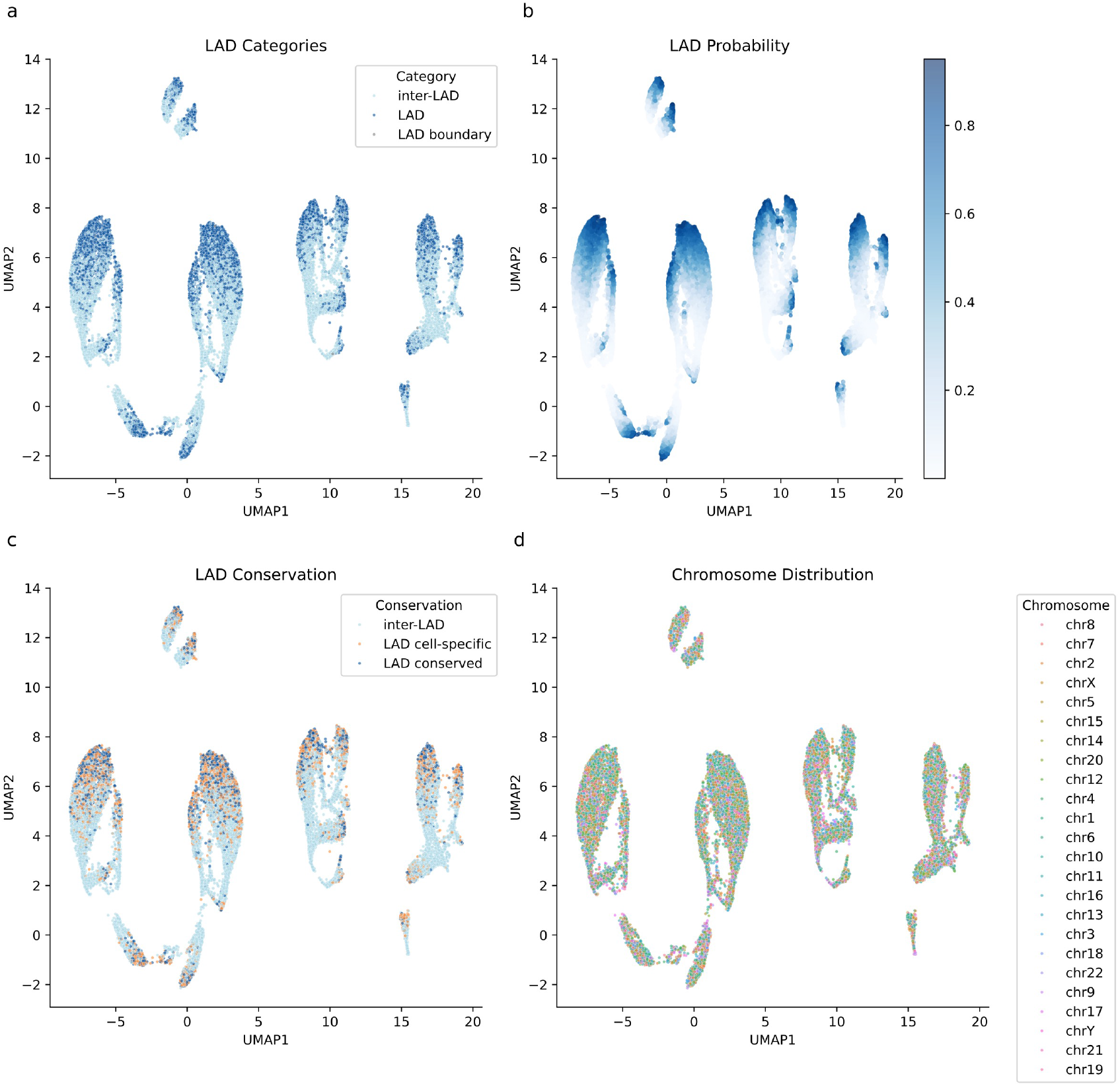
UMAP analysis of Nucleotide GPT’s learned sequence representations. Uniform Manifold Approximation and Projection (UMAP) visualization of genomic sequence embeddings from the final layer of Nucleotide GPT. (a) UMAP colored by LAD category shows distribution of LADs (blue), inter-LADs (light blue), and LAD boundaries (grey). (b) The same embeddings colored by LAD probability (>0.8, dark blue; <0.2, light blue) reveal strong correspondence between model predictions and actual LAD annotations. (c) When colored by LAD conservation status, conserved LADs (blue), RG-specific LADs (orange), and inter-LADs (light blue) show overlapping distributions in the embedding space, suggesting shared sequence features between constitutive and cell-type specific LADs despite differences in classification confidence. (d) Distribution of sequences by chromosome demonstrates no strong chromosome-specific biases in the learned representations.

When examining these same embeddings colored by LAD conservation status, we do not observe clear spatial segregation between conserved and RG-specific LADs (**Fig. 4c**). Instead, these two categories appear to occupy largely overlapping regions in the embedding space, suggesting that the sequence features learned by the model do not strongly distinguish between conserved and cell-type specific LADs.

### Nucleotide GPT Captures Nuclear Speckle Association Patterns

We also fine-tuned Nucleotide GPT to classify sequences associated with SPADs, inter-SPADs, and SPAD boundaries in the three cell types. For the RG sample, the model achieved 79.9% accuracy after 9,000 training steps. For the IPC sample, the model reached an accuracy of 80.1%. For the eN sample, the model achieved 79.7% accuracy. We also fine-tuned Nucleotide GPT on a combined dataset encompassing all three cell types to create a cross-cell type model. This model achieved an overall accuracy of 81.0% on SPAD classification.

We next examined how the model’s embeddings represent SPADs. Similar to our LAD analysis, we visualized the embeddings colored by the model’s ground-truth SPAD annotations (**Fig. 5a**) and predicted inter-SPAD probabilities (**Fig. 5b**). The UMAP visualization suggests that the model also learns to distinguish SPAD and inter-SPAD regions, with the predicted probabilities (shown in varying purple intensities) closely correspond to the actual SPAD classes (blue and orange annotations). As with LADs, the continuous nature of the probability distribution suggests the model captures a spectrum of nuclear speckle association strengths rather than binary classifications.

**Figure 5:**
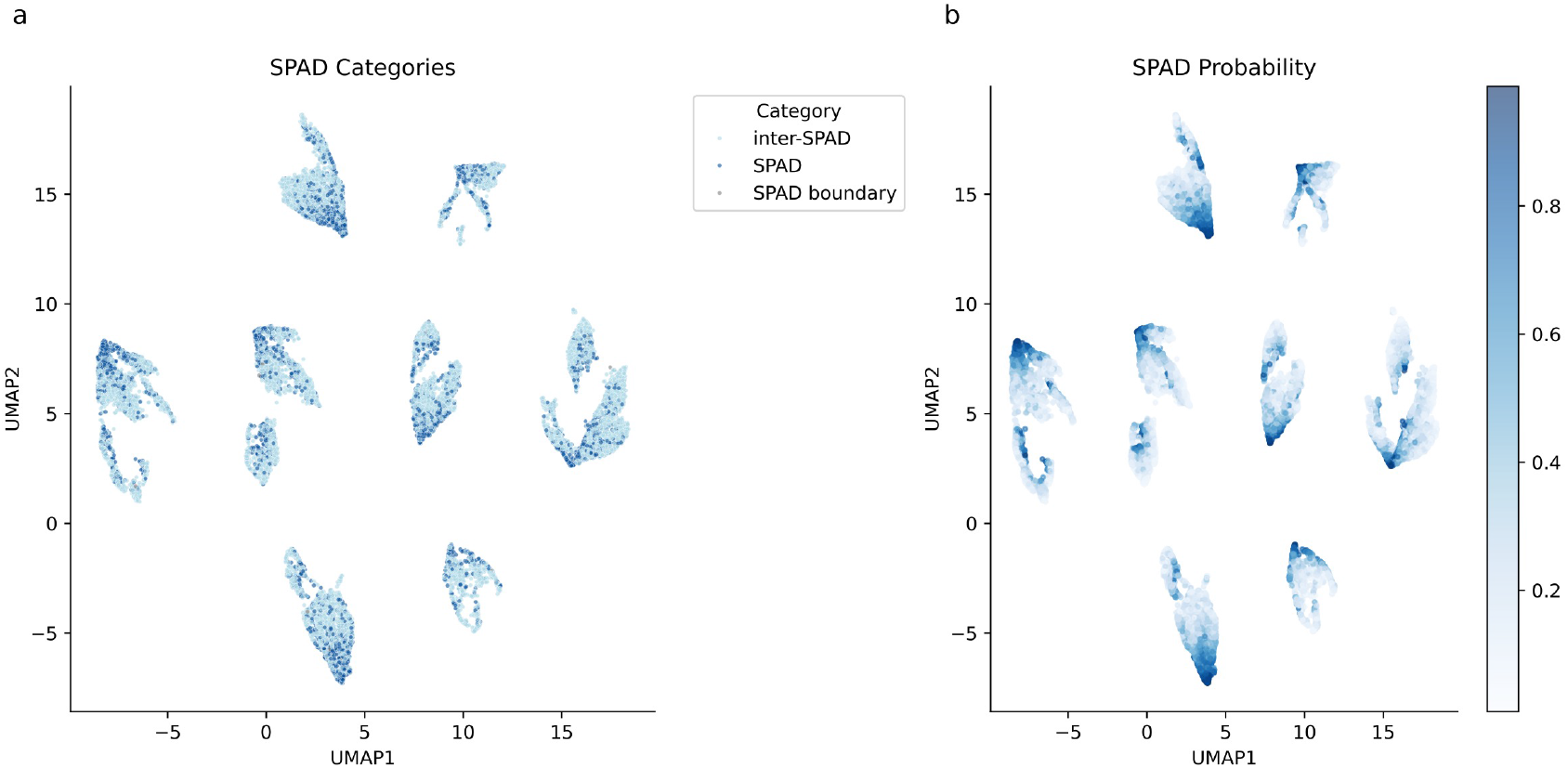
UMAP analysis of Nucleotide GPT’s learned sequence representations. UMAP visualization of genomic sequence embeddings from the final layer of Nucleotide GPT. (a) UMAP colored by SPAD category shows distribution of SPADs (blue), inter-SPADs (light blue), and SPAD boundaries (grey). (b) The same embeddings colored by SPAD probability (>0.8, dark blue; <0.2, light blue) reveal strong correspondence between model predictions and actual SPAD annotations.

### Nucleotide GPT Attention Patterns Provide Framework for Investigating Sequence Features Relevant to Classification

To better understand how our model makes decisions when classifying nuclear subcompartment associations, we developed a method to analyze the transformer’s attention patterns.

For each position in the DNA sequence, attention scores represent the cumulative importance assigned to that position by all attention heads across all layers of the model. By summing these attention scores, we created a single interpretable signal that can be visualized alongside other genomic annotations in genome browsers.

Preliminary exploration of these attention patterns suggests potentially interesting relationships with known genomic elements. **Fig. 6a** shows an example of attention patterns in a 7,300 bp region on chromosome 4 that the model correctly predicted to be a LAD in the eN sample. The attention track displays several attention spikes, where a spike is defined as a position in the sequence with an attention score more than three z-scores above the mean, including one notable peak with an attention score more than ten z-scores above the mean. The most notable peak overlaps with a region with high ReMap ChIP-seq density. In an RG LAD region on chromosome 3 (**Fig. 6b**), multiple attention peaks correlate with ReMap ChIP-seq signals and conserved genomic elements, suggesting the model recognizes these regulatory features as important for LAD classification. A constitutive LAD region on chromosome 10 (**Fig. 6c**) shows attention peaks that align with ReMap ChIP-seq signals and regulatory elements annotated by ENCODE and ORegAnno, indicating sequence features that may be important for maintaining stable LAD associations. Comparing attention patterns between different nuclear compartments reveals interesting contrasts. In a RG SPAD region on chromosome 2 (**Fig. 6d**), we observed attention peaks that specifically correspond to *ASXL2* exons and conserved genomic elements. This pattern differs notably from what we observed in a RG LAD region on chromosome 9 (**Fig. 6e**), where the model pays minimal attention to the *ADAMTSL1* exon. This differential attention to exonic regions between LADs and SPADs could indicate that the model has learned to distinguish these subcompartments partly based on their gene architectural features. These patterns collectively suggest that our model has learned to recognize specific sequence features associated with different nuclear subcompartments, with particular attention paid to regulatory elements, conservation patterns, and gene structure.

**Figure 6:**
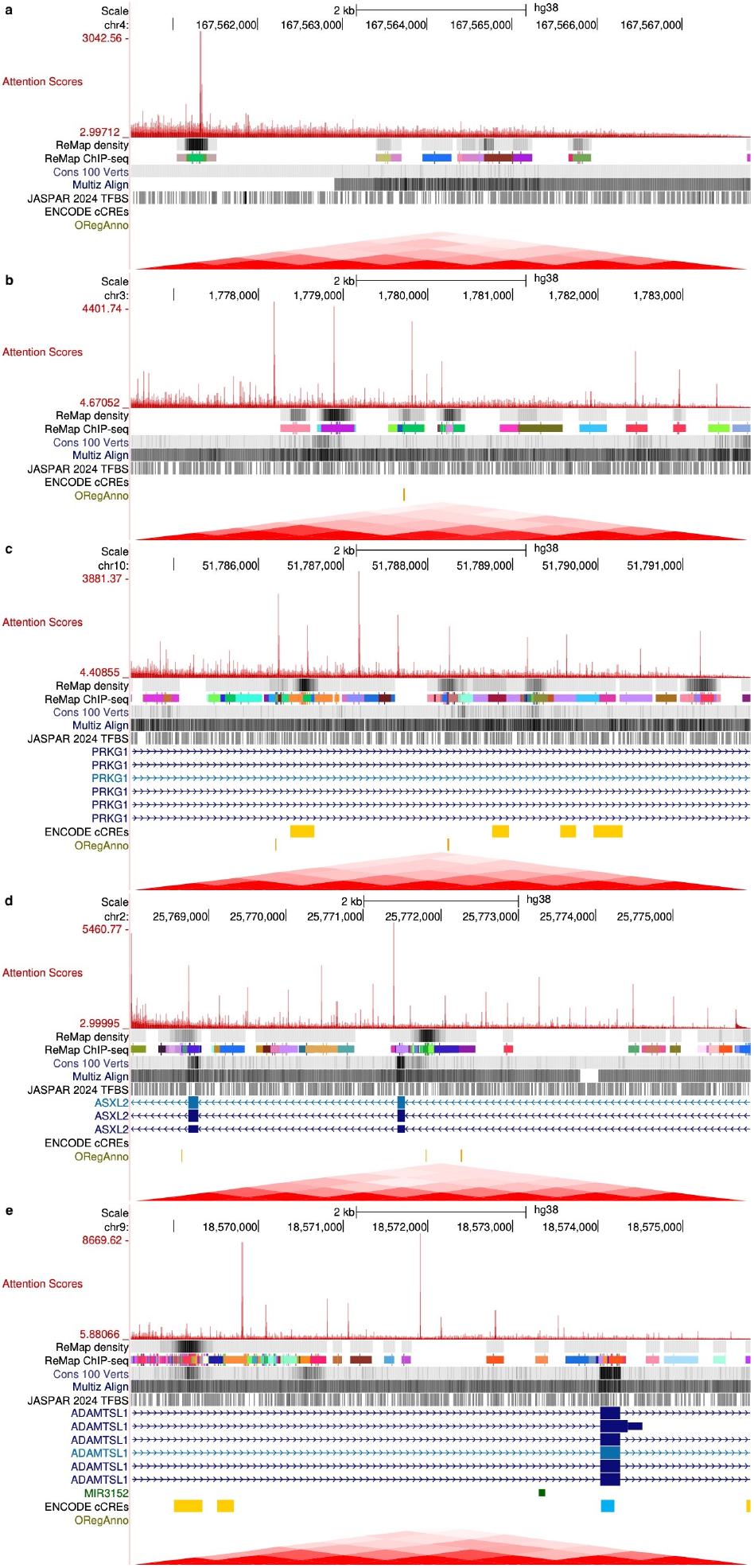
Transformer attention patterns reveal sequence features associated with nuclear compartment organization. Transformer attention scores (red), summed from all layers and heads, aligned with genomic annotation tracks, including ReMap ChIP-seq density, conservation scores (Cons 100 Verts and Multiz Align), JASPAR 2024 transcription factor binding sites, regulatory elements from ENCODE and ORegAnno databases, and Hi-C chromatin interaction data. The height of attention peaks indicates the relative importance assigned to specific sequence positions by the transformer model during classification.a) Genome browser view of chromosome 4 (chr4:167,560,510-167,567,809) showing attention scores (red) in an eN LAD region. A prominent attention spike coincides with an area of high ReMap ChIP-seq density, indicating dense transcription factor binding. Additional tracks show conservation scores (Cons 100 Verts, Multiz Align), JASPAR 2024 transcription factor binding sites, ENCODE candidate cis-regulatory elements (cCREs), and ORegAnno regulatory annotations. b) Genome browser view of chromosome 3 (chr3:1,776,001-1,784,000) displaying attention patterns in a RG LAD region. The attention track shows multiple distinct peaks correlating with ReMap ChIP-seq signals and conserved genomic elements, illustrating the model’s focus on regulatory features within LADs. c) Genome browser view of chromosome 10 (chr10:51,784,001-51,792,000) showing attention patterns in a constitutive LAD (cLAD) region, with peaks correlating with ReMap ChIP-seq signals, ENCODE cCREs and ORegAnno regulatory annotations. d) Genome browser view of chromosome 2 (chr2:25,768,001-25,776,000) depicting attention patterns in a RG SPAD region, with attention peaks corresponding to *ASXL2* exons and conserved genomic elements. e) Genome browser view of chromosome 9 (chr9:18,568,001-18,576,000) showing attention patterns in a RG LAD region. There is very little attention paid to the *ADAMTSL1* exon, in contrast to (d), which shows peaks aligned with exons.

## 4 Discussion

Nucleotide GPT demonstrates that transformer-based models can effectively learn complex patterns in genomic sequences and predict nuclear subcompartment associations with moderate accuracy. Our model achieved performance comparable to state-of-the-art genomic foundation models across diverse prediction tasks, including promoter detection, splice site identifications, and transcription factor binding site prediction.

A key consideration in applying transformer architectures to genomic sequences is how to tokenize DNA for model input. Unlike natural language, which has inherent structure through words and punctuation, DNA sequences consist of just four nucleotides without obvious segmentation. Most existing transformer-based genomic models operate on limited sequence lengths of 512 to 4,000 tokens, representing less than 0.001% of the human genome (22). Previous work on transformer-based prediction of spatial distance to nuclear subcompartments employed a k-mer based approach that reduces dimensionality by computing frequencies of fixed-length nucleotide patterns (16). While models like DNABERT, DNABERT-2, Nucleotide Transformer, and dnaGrinder use k-mer tokenization or byte-pair encoding to aggregate DNA sequences into larger units, Nucleotide GPT joins HyenaDNA in processing sequences at the individual nucleotide level (5) (4) (23) (24) (22). Unlike HyenaDNA which uses implicit convolutions to achieve longer sequence lengths, Nucleotide GPT employs a traditional transformer architecture with attention mechanisms, limiting sequence length but potentially enabling more flexible long-range interactions within its 8kb context window. This design choice reflects a tradeoff between sequence length capability and architecture complexity - while HyenaDNA can process sequences up to 1 million nucleotides, Nucleotide GPT’s 8kb window allows sufficient context for nuclear compartment prediction while maintaining the interpretability advantages of attention mechanisms.

Nucleotide GPT employs Flash Attention (20), an IO-aware attention algorithm that enables efficient training and inference while maintaining exact attention computations, alongside rotary positional embeddings (RoPE) (19) to encode positional information. Unlike previous genomic foundation models that use absolute positional encodings (5) or attention with linear biases (ALiBi) (35) (4) (24), RoPE directly encodes relative position information into the key-query attention computations through a rotation matrix. This approach has proven highly effective in state-of-the-art language models such as LLaMA and Grok (36). The choice of RoPE is particularly relevant for genomic sequences, where relative distances between sequence elements often carry more biological meaning than their absolute positions.

The model’s ability to predict LAD and SPAD associations with moderate accuracy (73.6% and 79.4% respectively) suggests that DNA sequence features play a significant role in determining nuclear organization. To the best of our knowledge, UNADON is the only other published model to test and demonstrate transformer-based prediction of nuclear subcompartment association (16). UNADON was developed using TSA-seq data from cultured cell lines to predict continuous cytological distance measurements to nuclear bodies. In contrast, Nucleotide GPT was trained on data from GO-CaRT, a method that maps nuclear compartment associations in tissue samples using CUT&RUN technology combined with fluorescence-activated nuclei sorting (FANS) (17) (34). The genome organization of cultured cells can differ substantially from those in vivo, with approximately 56% of previously annotated cell-type invariant LADs showing cell culture-specific associations.

By using GO-CaRT data from human brain tissue samples, Nucleotide GPT learns to predict discrete LAD and SPAD categories based on nuclear organization states present in vivo during cortical development. While our approach shares UNADON’s foundation in transformer architecture, the two models differ fundamentally in their input representations and architectural choices. UNADON integrates both DNA sequence features (through k-mer frequencies) and epigenomic signals as parallel inputs, using a multi-modal architecture to process these different data types independently before combining them. In contrast, Nucleotide GPT operates directly on raw DNA sequences at single-nucleotide resolution, demonstrating that sequence features alone contain sufficient information to predict nuclear compartment association. The models also differ significantly in their context handling. UNADON processes 5Mb regions but reduces sequence complexity through k-mer frequency vectors, while Nucleotide GPT maintains complete sequential information within an 8kb bp window.

LADs have been previously characterized by distinct genomic characteristics compared to inter-LADs, including low gene density, rich in long interspersed nuclear elements (LINEs), and high sequence A/T content (2). While many features that have been associated with LADs are epigenomic and context-dependent, the model’s performance indicates that underlying sequence composition and organization may play an important role in determining LAD formation. This aligns with previous work demonstrating that specific DNA sequences can direct chromatin association with the nuclear lamina. Zullo et al. (37) identified discrete lamina-associating sequences (LASs) enriched for GAGA motifs that are sufficient to establish lamina association and gene repression. Nucleotide GPT may be detecting these and other sequence patterns that facilitate LAD formation through interactions with proteins that facilitate chromatin tethering to the nuclear lamina.

While LAD prediction accuracy decreased from 77.5% in RG cells to 68.9% in eN cells, SPAD prediction remained relatively stable across cell types (80%). This pattern suggests that the sequence determinants of LAD associations may become more complex and/or context-dependent during neuronal differentiation. Recent work from our lab has shown that during cortical neurogenesis, the genome undergoes extensive remodeling of its interactions with nuclear compartments, with hundreds of neuronal genes relocating from the lamina to nuclear speckles (34). LAD architecture undergoes extensive remodeling, with approximately 19% of the genome changing its lamina association state during differentiation from RG to eN. The decreased accuracy of our sequence-based predictions in neurons might reflect this developmental complexity, suggesting that while sequence features provide a strong foundation for LAD formation in neural progenitors, additional cell-type specific factors likely modulate these associations during differentiation.

Our analysis of transformer attention patterns provides a novel window into how these models process genomic sequence information. By aggregating attention scores across layers and heads, we were able to identify specific genomic regions that the model consistently focuses on when making predictions about nuclear organization. Many of these attention spikes correspond to known regulatory elements and functionally important genomic features, with different patterns between LADs and SPADs. This suggests that the model has learned to recognize biologically relevant sequence patterns without explicit supervision over these features. The correspondence between attention spikes and known functional elements not only validates our approach but also suggests that attention score analysis could be used to identify which regulatory regions or sequence elements are important for LAD and SPAD architecture.

While our study demonstrates the potential of transformer models for predicting nuclear subcompartment associations, several limitations should be noted. Our single-nucleotide tokenization approach, while preserving complete sequence information, results in longer input sequences that precludes the model from capturing very long-range dependencies beyond the 8kb window size. This could affect the learning of higher-order organization features that span larger genomic distances. Previous deep learning approaches to predicting higher-order genome organization have relied on convolutional neural networks (CNNs), which have demonstrated the ability to process very long sequences (1Mb) through hierarchical architectures that progressively reduce dimensionality. For example, the Akita model uses a series of convolution and pooling operations to iteratively reduce 1Mb sequences to 2kb bins, enabling it to capture long-range regulatory interactions and domain-level genome organization (38). While our 8kb bp window size represents an improvement over previous models, it still constrains the model’s ability to capture very long-range genomic interactions that may influence nuclear organization.

This work represents a significant contribution to our understanding of the sequence determinants of nuclear organization. By demonstrating that transformer models can predict nuclear subcompartment associations from sequence alone, we provide additional evidence for the role of DNA sequence in determining 3D genome architecture. The model’s varying performance across cell types and its attention patterns provide new insights into how sequence features may contribute to both constitutive and cell-type specific aspects of nuclear organization. These findings lay the groundwork for future investigations into the mechanistic relationships between DNA sequence and nuclear architecture.

